# Response outcome gates the effect of spontaneous cortical state fluctuations on perceptual decisions

**DOI:** 10.1101/2021.09.01.458539

**Authors:** Davide Reato, Raphael Steinfeld, André Tacão-Monteiro, Alfonso Renart

## Abstract

Sensory responses of cortical neurons are more discriminable when evoked on a baseline of desynchronized spontaneous activity, but cortical desynchronization has not generally been associated with more accurate perceptual decisions. Here we show that mice perform more accurate auditory judgements when activity in the auditory cortex is elevated and desynchronized before stimulus onset, but only if the previous trial was an error, and that this relationship is occluded if previous outcome is ignored. We confirmed that the outcome-dependent effect of brain state on performance is neither due to idiosyncratic associations between the slow components of either signal, nor to the existence of specific cortical states evident only after errors. Instead, errors appear to gate the effect of cortical state fluctuations on discrimination accuracy. Neither facial movements nor pupil size during the baseline were associated with accuracy, but they were predictive of measures of responsivity, such as the probability of not responding to the stimulus or of responding prematurely. These results suggest that the functional role of cortical state on behavior is dynamic and constantly regulated by performance monitoring systems.

---

Successfully performing any behavior, including the acquisition and processing of sensory information to guide subsequent action, requires that the dynamical regimes of neural circuits across the whole brain be set appropriately in a coordinated fashion. The activation-inactivation continuum – the degree to which the activity of cortical neurons tends to fluctuate synchronously and in phase on time-scales of hundreds of milliseconds^1–3^ – and pupil dilation – a measure of cognitive load and arousal^4,5^ – are commonly used to label these large-scale dynamical regimes, often referred to as ‘brain states’^6–11^. What is the relationship between cortical state and behavior? Although cortical desynchronization during wakefulness in rodents was initially linked to movement during exploration^3^, desynchronization and movement can be dissociated^9,11^. In fact, it was demonstrated early that desynchronization can occur under immobility during visual attention^12,13^, suggesting that desynchronization during waking might signal a state where the animal’s cognition is oriented towards the environment^3^. Such state would presumably be associated with the ability to perform finer perceptual judgments – a hypothesis consistent with many studies showing that the discriminability of neural sensory representations increases monotonically with the level of cortical desynchronization^14–18^. Behavioral studies, however, have not generally confirmed this picture. During sensory detection, performance and arousal (which tends to be associated with desynchronization^9,10^) are related, but in a non-monotonic fashion^10,19^. Furthermore, detection tasks are limited in their ability to decouple sensory discrimination and the tendency of the subject to respond, which is relevant since both aspects are potentially associated with changes in brain state. Two-alternative forced-choice (2AFC) discrimination tasks allow a cleaner separation between responsivity and accuracy, but two studies using this approach failed to find a clear link between desynchronization and perceptual accuracy, pointing instead to a role on task engagement, responsivity and bias^20,21^(but see^17^ for effects of desynchronization during the delay period of a delayed comparison task). Thus, existing evidence suggests that the effects of desynchronization on discriminability at the neural and behavioral levels are not fully consistent, raising questions about the functional role of the desynchronized state. Here we suggest a possible explanation for this discrepancy, by showing that the effect of desynchronization on accuracy during an auditory 2AFC discrimination task depends strongly on the outcome of the previous trial, and is occluded if trial outcome is ignored.

## Movement, arousal and temporal fluctuations in baseline activity

In order to investigate the impact of cortical desynchronization on discrimination accuracy, we recorded population activity from the auditory cortex (Supplementary Fig. 1a) of head-fixed mice while they performed a 2AFC delayed frequency discrimination task (Fig. 1a-c, Methods). In addition, we monitored pupil size (PupilS) as well as the overall optic flow (OpticF; Fig. 1d, Methods) of a video recording of the face of the mouse (Supplementary Fig. 1b), as a proxy for movement signals known to affect synchronization^3,8,22^ and cortical activity^23–25^.

**Fig 1.**
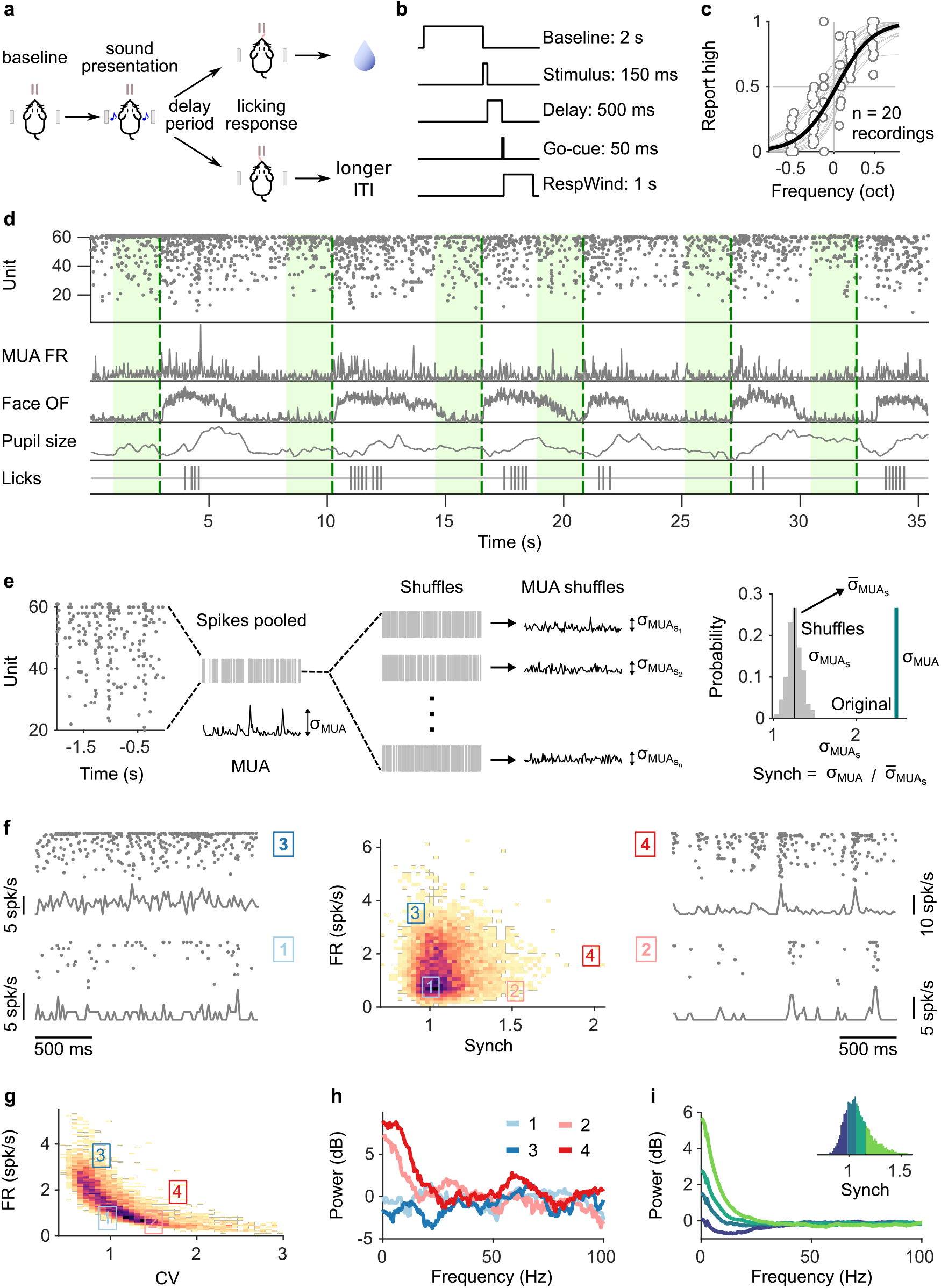
Task structure, signals monitored, and quantification of synchrony in baseline activity. **(a)** Task schematic. Head-fixed mice lick at one of two spouts depending on whether the frequency of a pure tone is higher or lower than 14 kHz. **(b)** Temporal sequence of events in a trial. Mice should respond after a delay of 0.5 s. Baseline activity is analyzed in a window of 2 s before the presentation of the sound. **(c)** Discrimination performance. Each dot is the proportion of times a mouse reports high in a given recording session to a given sound. Solid curve is a logistic regression fit. **(d)** Signals monitored. Top to bottom are population raster, multiunit firing rate (MUA FR), mean optic flow of the face (OpticF), size of the pupil (PupilS) and licks. Dashed vertical lines mark stimulus presentation times and green background marks the baseline period we analyze. **(e)** Method for quantifying synchronization. Synch effectively measures the population averaged correlation in the baseline period relative to surrogate data with the same number of spikes but randomly placed in the same period of time (Methods). **(f)** Distribution of baseline FR and Synch pooled across all recording sessions. Plots on the sides show rasters and population firing rates for four example baseline periods. **(g)** Identical plot to the one in (f)-middle, but where global synchronization is assessed using the coefficient of variation (CV) of the instantaneous population rate (Methods). CV and FR are negatively correlated. **(h)** Power spectrum (Methods) of the four individual example baseline periods in (f). **(i)** Average power spectrum of each of the four quantiles of the distribution of Synch across trials. Large values of Synch reflect low-frequency coordinated fluctuations across the population. Inset: aggregate distribution of Synch values across recordings. Each quantile corresponds to one of the spectra in panel (i).

The dynamical regime of baseline spontaneous activity in the auditory cortex in a period of 2 s prior to the presentation of the stimulus was quantified using two statistics: overall firing rate (FR) across the population, and degree of synchronization (Synch). In order to obtain a measure of synchronization as independent as possible of FR, we quantified Synch for each baseline period relative to surrogates of the spike trains from the same period (thus with equal surrogate FR) but shuffled spike times (Fig. 1e,f, Methods). This measure is normalized, and would take a value of 1 if neurons were statistically uncorrelated and displayed Poisson-like firing. We found that the resulting Synch and FR measures were effectively uncorrelated (Fig. 1f), to a much larger extent than previously used measures of synchronization, such as the coefficient of variation of the multiunit activity^18,26^ (Methods), which displayed negative correlations with baseline FR (Fig. 1g). The coordinated fluctuations responsible for Synch are of low frequency, as evident from trial-to-trial comparison of Synch and the power-spectral density of the MUA (Fig. 1h,i; Methods). In particular, strong desynchronization was associated to a suppression of power in the ∼ 4 – 16 Hz frequency band relative to a Poisson spike train of the same firing rate (Fig. 1i).

Before inspecting the relationship between each of these four signals (OpticF, PupilS, FR and Synch) and discrimination performance, we explored the way in which PupilS and OpticF shape baseline neural activity. To do this, we separately regressed FR and Synch on PupilS and OpticF using a linear mixed model with recording session as a random effect (Methods). This analysis revealed FR to be associated to movement and pupil size (Fig. 2a, top). Surprisingly, Synch did not show a clear association with either predictor (Fig. 2a, bottom), and a tendency to increase with pupil size, contrary to previous findings^9,11^. Seeking to understand this puzzling result, we inspected more carefully the time-series for each of the four baseline signals. This revealed that, in addition to fast trial-by-trial fluctuations, there exist both clear session trends as well as slow fluctuations spanning many trials, leading to broad auto- and cross-correlations (Supplementary Fig. 2). These slow components – presumably determined by slow physiological processes^27^which we don’t control – generically lead to correlations between the signals even if the trial-by-trial fluctuations that we are interested in are independent^28–31^. To address this problem, instead of using the raw signals, we focused on their innovations, i.e., that aspect of each signal which could not be predicted from itself and all others in the past (Fig. 2b). Formally, we defined the innovation associated to each signal (which we denote with the subscript I, e.g., FR_I_) as the residual of a multivariate linear fit of the raw signal using as predictors its own value and that of the other 3 signals up to 10 trials into the past, the trial number in the session to account for slow within-session trends, and the outcome (correct/incorrect) of the 10 previous trials (Supplementary Fig. 3a; Methods). Different signals could be predicted by past information to different extents, with Synch and PupilS being the least and most predictable respectively (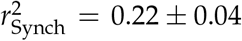 and 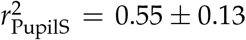 median ± MAD across recordings; Supplementary Fig. 3b,c). Innovations, on the other hand, displayed effectively ‘white’ auto- and cross-correlations (Fig. 2c). Thus, any associations revealed using innovations as regressors will not be caused by idiosyncratic associations between slowly varying processes correlated with the raw baseline signals^28–31^. When the analysis in Fig. 2a was repeated using innovations, a different picture emerged. Although FR_I_ is positively correlated with both OpticF_I_ and PupilS_I_ (Fig. 2c), the correlation with PupilS_I_ is explained away by the positive correlation between OpticF_I_ and PupilS_I_ themselves, revealing a clear positive association only between movement and FR innovations during the baseline (*p* < 0.0002, bootstrap quantile method^32^, from now on referred to as ‘bootstrap’; Methods). Synch_I_ is much more weakly correlated with both OpticF_I_ and PupilS_I_ (Fig. 2c,d). Nevertheless, the analysis revealed a positive association between pupil size and desynchronization (*p* < 0.0002, bootstrap) – consistent with previous studies^9,11^ – as well as a rather small but significant (*p* = 0.012, bootstrap) positive association between movement and synchronization (Fig. 2d). For the rest of our study, we seek to explain choice behavior in terms of innovations to characterize trial-by-trial relationships between discrimination accuracy and brain state uncontaminated by slow physiological trends.

**Fig 2.**
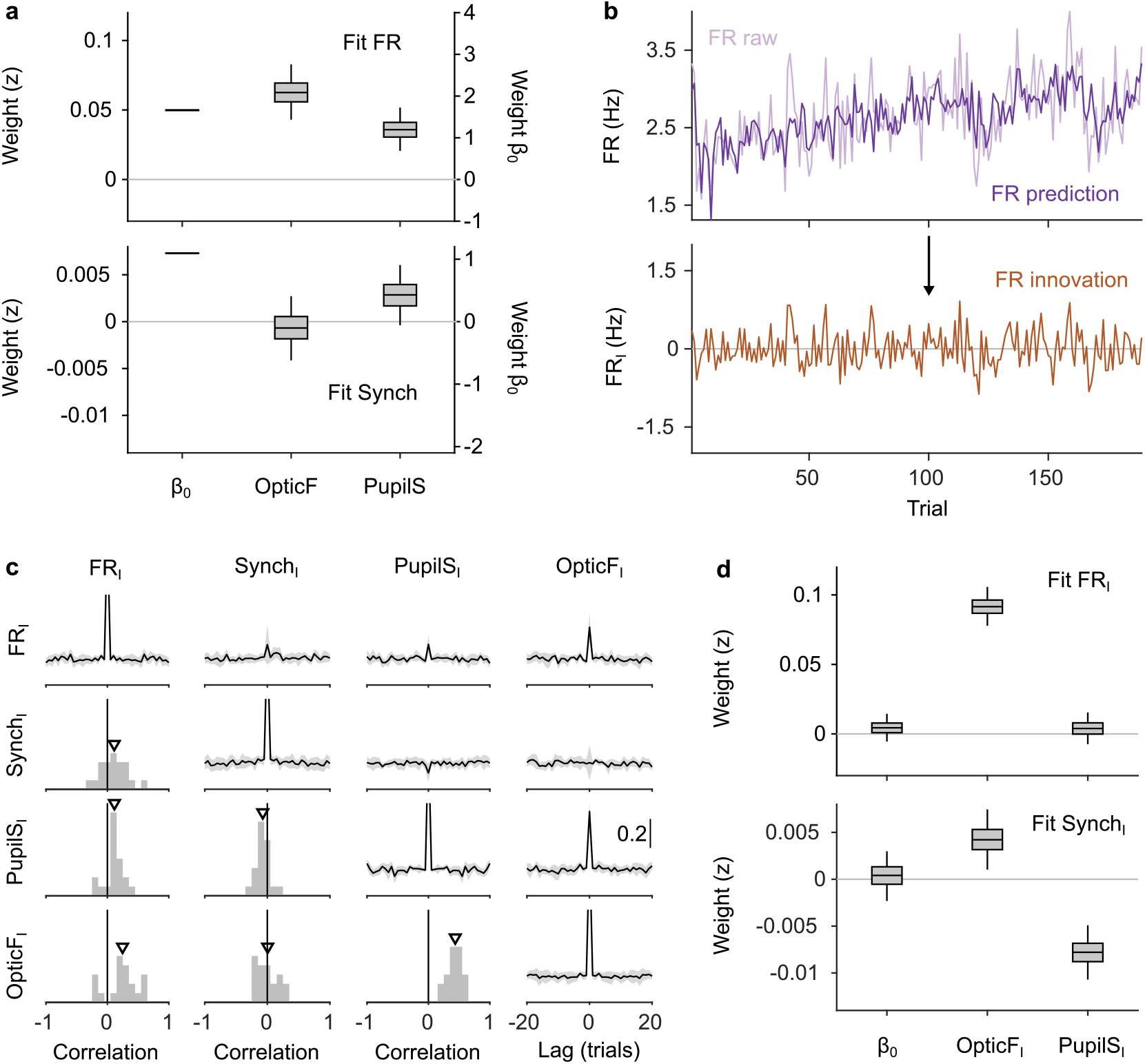
Innovations clarify the effect of movement and pupil size on cortical state fluctuations. **(a)** Linear mixed model regression (Methods) of FR (top) and Synch (bottom) on movement and pupil size. Graphs show values of regression coefficients. Box plots here and elsewhere represent median, interquartile range and 95% CI on the bootstrap distribution of the corresponding parameter (Methods). Offset can be read from the right y-axis. **(b)** Example of the process of calculating innovations for the baseline FR of one recording session. Top, raw data and prediction of the raw data (Supplementary Fig. 3; Methods). The innovation FR_I_ (bottom) is the difference (prediction residual) between the two traces in the top. **(c)** Correlation between OpticF, PupilS, FR and Synch innovations. Diagonal and above, cross-correlations between each of the four signals (black, median across recordings; gray, median absolute deivation (MAD)). Below diagonal. For each pair of innovations, histogram across recordings of their instantaneous correlation. Triangles mark the median across recordings. **(d)** Identical analysis as panel (a) but using innovations instead of the raw signals.

## Outcome-dependent effect of desynchronization on choice accuracy

We used a generalized linear mixed model (GLMM; Methods) to explain whether each trial was correct or an error based on the strength of sensory evidence (Stim) and the four innovations during the baseline preceding that trial. We sought to predict whether a choice was correct rather than the choice itself (left versus right) so that the potential effect of innovations would represent a main effect in the model, rather than an interaction with the stimulus (but see Supplementary Fig. 4a). This analysis only considers valid trials (Methods) where the mice made a choice within the response window, and thus quantifies the effect of brain state on discrimination accuracy regardless of unspecific response tendencies. In order to be able to explain within-session trends, we always include a regressor coding the trial number within the session (TrN). Finally, to model possible sequential dependencies in choice-accuracy, we also included a regressor with the outcome (correct/error) of the previous trial (pCorr; only valid previous trials were considered). The analysis revealed a positive association between TrN and accuracy (Fig. 3a; *p* = 0.005, bootstrap) – reflecting the fact that mice tend to become more accurate throughout the session – but none of the four baseline predictors had an association with accuracy^21^ (Fig. 3a; Supplementary Table 1 lists the complete results of all GLMM fits in the main text). However, the coefficient measuring the effect of the outcome of the previous trial was negative (*p* = 0.006, bootstrap), suggesting that mice tended to be more accurate after errors (Fig. 3a). Indeed, across sessions, accuracy was larger after an error (Fig. 3b; *p* =0.021, signrank test, Methods). It is well known that errors have an effect on the reaction time (RT) of the subsequent trial^33–35^, and, although less consistently, accuracy enhancements after an error have also been observed^34–36^. Given that errors have an impact on task performance, we reasoned that they might modulate the role of spontaneous cortical fluctuations on choice. To test this hypothesis, we performed our analysis separately after correct and error trials.

**Fig 3.**
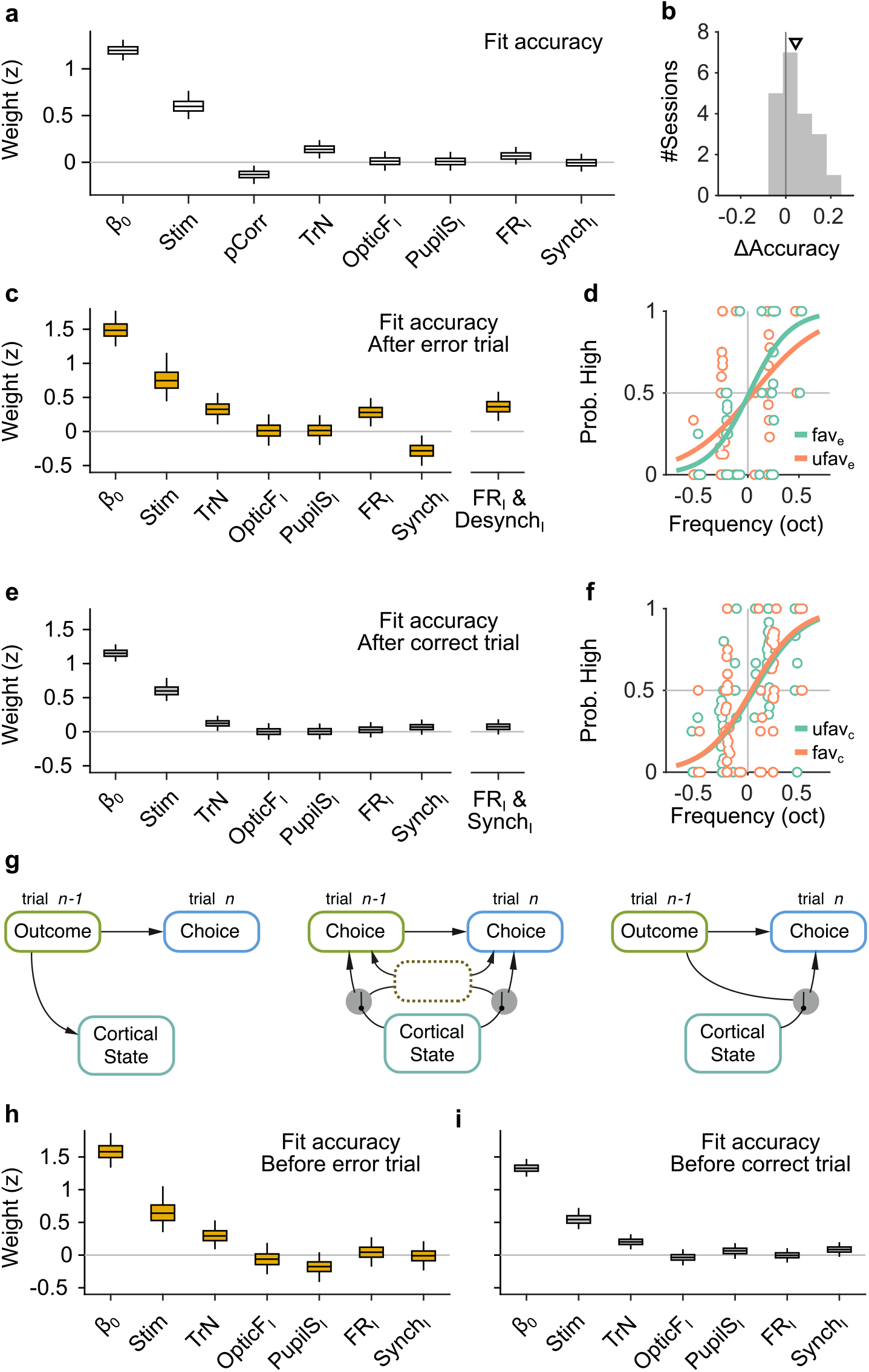
The effect of spontaneous state fluctuations on accuracy is outcome dependent. **(a)** Coefficients of a GLMM fit to the mice choice accuracy in valid trials. Accuracy is affected by the strength of evidence, the point during the session and the outcome of the previous trial, but none of the four signals computed during the baseline explain accuracy. **(b)** Mean difference in accuracy after errors minus after corrects in each of the recording sessions. Triangle, median across sessions. **(c)** GLMM fit to accuracy computed separately after error trials. On the right we show the distribution of a single coefficient capturing trial to trial fluctuations in desynchronization and firing rate simultaneously (see text). **(d)** Psychometric function (logistic fit, Methods) of aggregate data across sessions separately for trials with favorable (Synch_I_(*z*) < 0 and FR_I_(*z*) > 0) and unfavorable (Synch_I_(*z*) > 0 and FR_I_(*z*) < 0) baseline states after a error trials. **(e**,**f)** Same as (c,d) but for choices after a correct trial. Note that, based on the results in (e), the favorable state after a correct trial is Synch_I_(*z*) > 0 and FR_I_(*z*) > 0. **(g)** Schematic illustration of possible relationships between outcome, baseline cortical state and accuracy. Left, the association between state and accuracy is spurious and results from a common effect of response outcome on these two variables. Middle, epoch hypothesis (see text). An unmeasured variable with a time-scale of several trials mediates both the effect of state on accuracy and the prevalence of errors. Right, response outcome gates the effect of state fluctuations (errors open the gate) on choice accuracy. **(h**,**i)** Same as (c,e) but conditioned on the outcome of the next, rather than the previous trial.

The results revealed that, while pupil size and movement still had no association with accuracy for either outcome separately (Fig. 3c,e), the effect of baseline neural activity on choice accuracy was indeed outcome dependent (Fig. 3c-f). After errors, both FR and Synch innovations in the baseline period explain accuracy (Fig. 3c; *p* = 0.0056 and *p* = 0.0124 for FR_I_ and Synch_I_ respectively; bootstrap). Mice made more accurate decisions when the baseline activity was higher and more desynchronized, a state we refer to as ‘favorable’ for accuracy after an error. In contrast, baseline activity had no clear association to accuracy after correct trials (Fig. 3e; *p* = 0.64 and *p* = 0.22 for FR_I_ and Synch_I_ respectively; bootstrap), despite the fact that the GLMM for after-correct choices had approximately three times as many trials (which is reflected on the smaller magnitude of the confidence intervals for this model; Fig. 3e). Although this makes it difficult to define a ‘favorable’ state for accuracy after correct trials, the median value of the coefficients for both FR_I_ and Synch_I_ in Fig. 3e is positive, suggesting that, if anything, more accurate choices after a correct trial were preceded by more synchronized (and stronger) baseline activity. The lack of effect of baseline activity on accuracy unconditional on outcome (Fig. 3a) is explained partly by the tendency of baseline fluctuations preceding a correct choice to have different signs (relative to the mean) after correct and error trials and by the fact that most trials (77%) are correct.

To assess together the effect of baseline FR and Synch innovations on accuracy, we created a single predictor for each baseline period whose value was equal to the projection of the (z-scored) two-dimensional pair (Synch_I_, FR_I_) onto a line of slope -45 deg on this plane (after errors), or 45 deg after corrects (Methods). This single predictor takes large positive values when both FR and Synch are ‘favorable’ for accuracy for each separate outcome. After errors, the combined effect of FR and Synch was 28% stronger than that of either of them separately and highly significant (Fig. 3c, rightmost coefficient; *p* = 0.0006, bootstrap), but it was still not significant after correct choices (Fig. 3e, rightmost coefficient; *p* = 0.2, bootstrap). To more directly quantify the effect of baseline neural activity on accuracy, we also computed aggregate psychometric functions for trials where the state of the baseline was favorable or unfavorable, separately after correct and error trials. The slope of the psychometric function was 68% larger in a favorable baseline (Synch_I_(*z*) < 0 and FR_I_(*z*) > 0) after errors (Fig. 3d; *p* = 0.04, permutation test, Methods). There was no visible effect of a favorable state after a correct trial (during which the cortex was more synchronized) on the aggregate psychometric function (Fig. 3f, *p* = 0.88, permutation test). Finally, we tested the robustness of these findings. The outcome dependence of the effect of baseline fluctuations on accuracy was qualitatively similar in model fits of trial-by-trial choice (as opposed to accuracy; Supplementary Fig. 4a, Methods), and also held when using parametric methods for the calculation of confidence intervals (Supplementary Fig. 4b, Methods).

What exactly do the results in Fig. 3a-f imply for the relationship between spontaneous baseline activity and choice? An explanation of these results as a spurious correlation caused by the joint influence of the outcome of the previous trial on accuracy and on baseline activity in the current trial (Fig. 3g, left) can be ruled out, since the outcome of the previous trial is fixed in the analyses of Fig. 3c,e. Rather, our results suggest that errors gate, or enable, the influence of spontaneous fluctuations on choice (Fig. 3g, right). However, it is still possible the gating is not performed by errors *per se*, but rather by some other quantity that tends to covary in time with errors. In other words, there might be epochs within the session during which spontaneous cortical fluctuations have an effect on accuracy and during which errors are more frequent (Fig. 3g, middle). We refer to this as the ‘epoch hypothesis’. The epoch hypothesis can be tested under the assumption that the epochs last a few trials, in which case the relationship between baseline activity and accuracy should be approximately symmetric around the time of an error. To test if this is the case, we repeated the analysis in Fig. 3c,e, but instead of conditioning on the outcome of the previous trial, we conditioned on the outcome of the next trial. If the epoch hypothesis is true, we would expect for FR_I_ and Synch_I_ to explain accuracy in a trial when the next trial is an error, just like in Fig. 3c. In contrast, we found that baseline fluctuations have no predictive power on the accuracy of a trial regardless of the outcome of the next trial (Fig. 3h,i). If trial *n* + 1 is correct, the influence of Synch_I_ and FR_I_ on the accuracy in trials *n* is similar to that observed if trial *n* – 1 is correct: not significantly different from zero but with a tendency towards higher accuracy when the baseline is more synchronized (Fig. 3e and Fig. 3h bottom). In contrast, baseline activity is clearly predictive of choice accuracy in trial *n* only if an error takes place in trial *n* – 1, but not on trial *n* + 1. These results are inconsistent with the idea that errors mark epochs of high correlation between cortical fluctuations and accuracy, and support instead the hypothesis that this correlation is triggered by the errors themselves (Fig. 3g, right).

Our analysis for the quantification of innovations in cortical state fluctuations revealed that there are slow time-scales associated to these signals (Supplementary Fig. 2) which, by construction, do not contribute to the effects in Fig. 3. Although the interpretation of possible correlations between mouse behavior and these slow signals is complicated^28,30,31^, we nevertheless assessed whether they were salient in our dataset. To do this, we smoothed, linearly detrended and z-scored the raw baseline FR and Synch time-series and the corresponding accuracy in those trials (Supplementary Fig. 5a, Methods), and computed their cross-correlation. We observed no correlations between Synch and accuracy (Supplementary Fig. 5b; *p* =0.94, signrank test) and a trend towards epochs of high performance to precede epochs of low baseline FR (Supplementary Fig. 5b; *p* =0.1, signrank test). Thus, even at slow time-scales, cortical state and decision accuracy are not saliently correlated, consistent with previous studies^21^ and with the outcome-independent analysis in Fig. 3a.

## Cortical fluctuations are only weakly affected by trial outcome

We next sought to understand whether the selective influence of baseline activity on choice after errors (Fig. 3) is due to a particular pattern of cortical state fluctuations that is only evident after the mouse makes a mistake. For instance, it’s possible that desynchronization is always conducive to better performance, but that sufficient levels of desynchronization are only attained after errors. We explored this question by quantifying the extent to which trial outcome shapes cortical state fluctuations (Fig. 4a). To accomplish this, we modified the definition of innovations for this set of analyses, excluding the outcome of the previous trials from the set of regressors that we use to compute the residuals (Methods). The values of Synch_I_ and FR_I_ observed after an error or a correct trial largely overlap (Fig. 4b), and the joint distribution of Synch_I_ and FR_I_ across recordings are very similar (Fig. 4c). We first quantified these effects calculating the signed discriminability index *d*′ (correct minus errors) of the distributions of Synch_I_ and FR_I_ for each recording. Across recordings, neither of these two measures were significantly different from zero (*p* = 0.26 and *p* = 0.13 for Synch_I_ and FR_I_ respectively; signrank test). As an alternative, more sensitive approach to understand which features of the baseline contained information about the outcome of the previous trial, we used a GLMM to decode whether the outcome of trial *n* – 1 was correct, using as regressors the four innovations in the baseline of trial *n*, as well as the session trend TrN. Previous-trial outcome is best explained by the PupilS_I_ in the subsequent baseline (Fig. 4e). This is intuitively clear, as correct trials are followed by licking, which is associated to pupil dilation^37^, a relationship that becomes obvious when plotting the cross-correlation function between the accuracy and PupilS time series (Fig. 4f). In addition to the pupil size, FR_I_ is also affected by the outcome of the previous trial, being smaller than average after correct trials (consistent with the small negative median value of 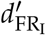 in Fig. 4d and with the negative trend in Supplementary Fig. 5b). Synch_I_ could not be used to predict the outcome of the previous trial. Overall, these results are not consistent with the effects in Fig. 3 being due to the presence of unique values of FR and Synch exclusively after errors. Errors do increase the FR in the next baseline period, but FR distributions after the two outcomes are largely overlapping. In addition, and somewhat unexpectedly, trial outcome has no effect at all on baseline synchrony.

**Fig 4.**
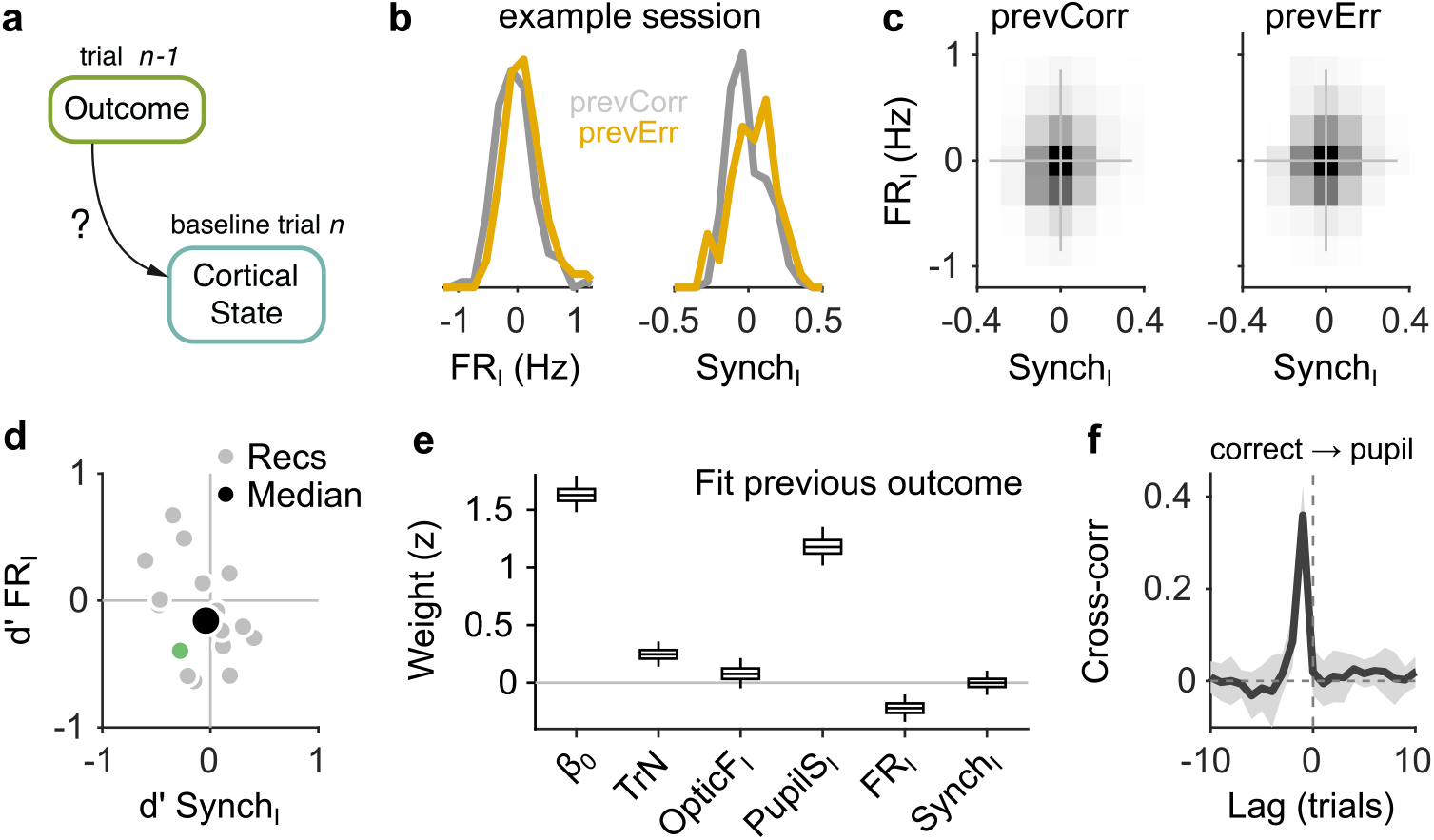
Effect of outcome on baseline activity. **(a)** Schematic illustration of the question addressed in this figure. **(b)** Distribution of FR_I_ (left) and Synch_I_ (right) after each of the two outcomes for an example session. **(c)** Joint histogram of FR_I_ and Synch_I_ on aggregate across recordings after a correct (left) and after an error (right) trial. **(d)** Discriminability index *d*′ between the distributions of FR_I_ and Synch_I_ (such as those in (b)) after each of the two outcomes. Each gray dot corresponds to one recording, the colored dot is the example recording in (b), and the large black circle is the median. **(e)** Coefficients of a GLMM fit to the outcome (correct or error) of the mice’s choices on trial *n* -1 using as regressors TrN and innovations from the baseline of trial *n*. **(f)** Cross-correlation function between the raw outcome and PupilS time series (Methods). Black is the median across recordings, gray is the MAD. Throughout this figure, innovations were modified so as to exclude previous outcomes in the calculations of the residuals (Methods).

## Effect of spontaneous fluctuations on measures of responsivity

Arousal and desynchronization have been shown to modulate measures of responsivity^10,20,21^. There are two different facets to responsivity in a discrimination task. One relates to the tendency of the subject to respond at all to a presented stimulus, which can be taken as a measure of task engagement. The other is RT, the time (since stimulus onset) it takes for the subject to respond. In a delayed response task like ours, there is additionally the possibility for mice to respond prematurely, failing to wait for the go signal at the end of the delay period (Fig. 5a). In our task, most trials were valid (either correct or errors, 70%, Methods), but there were also premature trials (7%) and ‘skips’ where the mice did not respond (23%; Fig. 5b).

**Fig 5.**
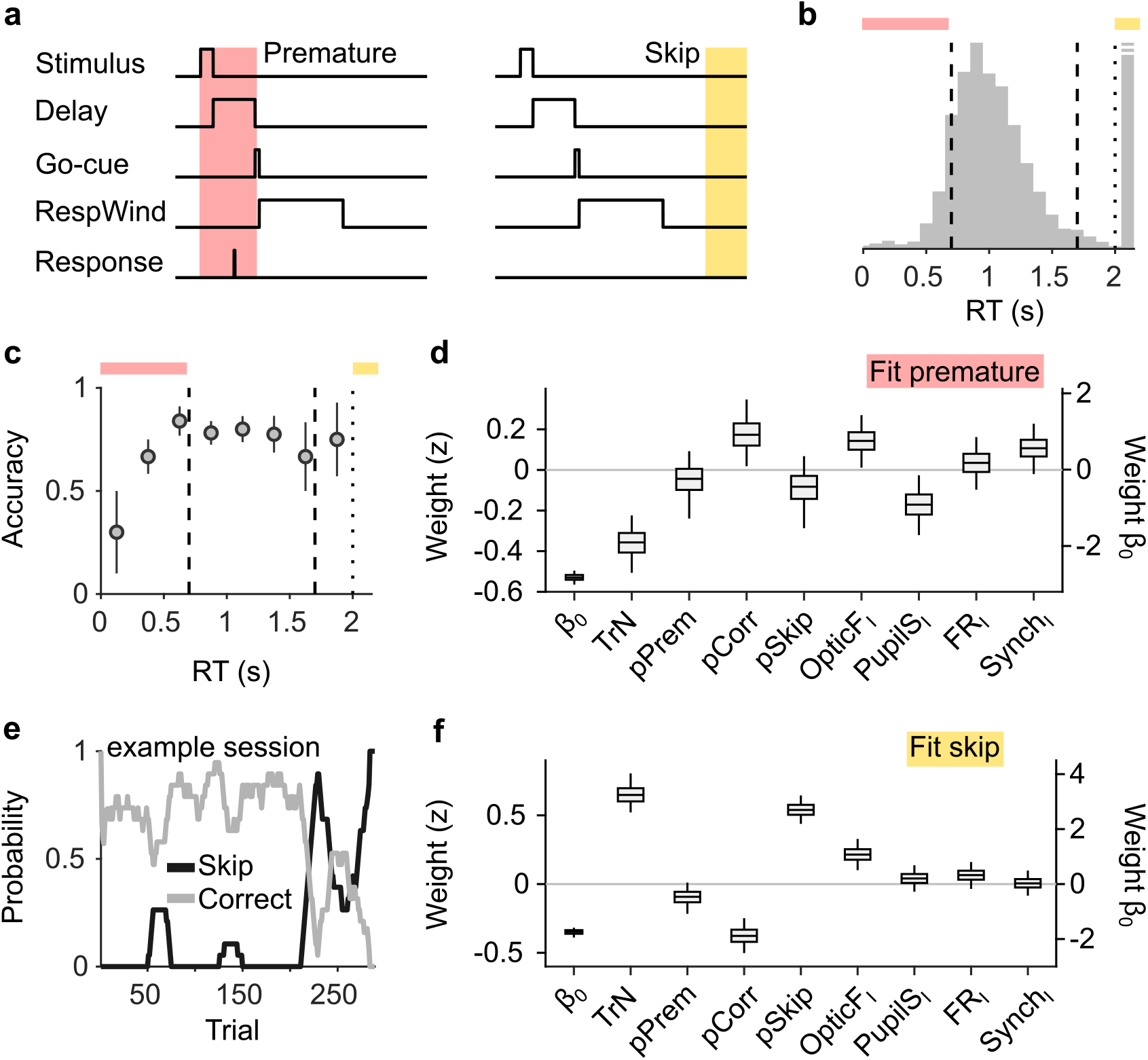
Effect of cortical state fluctuations on premature responding and engagement. **(a)** Definition of premature responses and skips. **(b)** Aggregate across sessions of the distribution of RTs in our task. Dashed lines indicate the response window in which a correct response was rewarded (valid trials). Trials where a response is not produced before the dotted line are defined as skips. Top, colors used to signal each trial type in (b). **(c)** Accuracy (median ± MAD across recordings) conditional on RT. **(d)** Coefficients of a GLMM fit to explain whether a given trial is premature or valid. Magnitude of the offset (*β*_0_) should be read of from the right y-axis **(e)** Probability of not responding to the stimulus (skip) in an example session. Skips tend to occur in bouts and are more frequent towards the end of the session. **(f)** Same as (d) but for a GLMM aimed at explaining if a particular trial is a skip or valid.

Choice accuracy varied as a function of RT (Fig. 5c). Very premature responses where most inaccurate. Accuracy tended to increase with RT for premature responses during the delay period, and then remained approximately constant within the valid response window and beyond. These results suggest that premature and valid responses might be differentially regulated. We explored this possibility by trying to explain whether a trial would be premature or valid using a GLMM. Unlike Figs. 3 and 4, which only deal with transitions between valid trials, here the previous trial could be either valid, premature or a skip, and we thus included corresponding regressors in the GLMM (Methods). The most reliable predictor of a premature trial was TrN (Fig. 5d; *p* < 0.0002, bootstrap), signalling a decreasing tendency to respond prematurely as the session progresses, paralleling changes in motivational state^38^. Everything else kept equal, premature trials also happened more frequently after correct trials (*p* = 0.03, bootstrap), in the presence of movement in the baseline (*p* = 0.03, bootstrap), and when cortical activity was more synchronized, although this last effect did not reach significance (*p* = 0.09, bootstrap). Interestingly, baseline periods with contracted pupil were predictive of premature responses (*p* = 0.02, bootstrap). Although this finding might seem at odds with previously reported associations between states of dilated pupil and impulsivity^10,21^, a large body of work has linked pupil dilation with the ability to exercise inhibitory control^39,40^, which is needed in order to avoid responding prematurely (Discussion).

Although RT did not primarily reflect decision time and was instead constrained by the delay period of the task (Fig. 1a,b; Fig. 5a), we nevertheless used a similar approach to explore a possible effect of cortical desynchronization on RT. Only movement innovations were positively associated with RT (*p* = 0.01, bootstrap; Supplementary Fig. 6). Somewhat surprisingly, we observed evidence of post-error slowing^33–35^, suggesting that the connection between errors and subsequent RT is so strong that it survives the constraints in RT imposed by a delayed response task.

Finally, we examined engagement. As commonly observed, mice underwent periods of disengagement during behavioral sessions^21,41^, defined as bouts of consecutive trials during which the mice didn’t respond to the stimuli (‘skips’, Fig. 5a,e). We attempted to predict whether a trial would be a skip or valid using identical regressors as for premature responses. Opposite to premature trials, skips were more frequent at the end of the session (Fig. 5f, *p* < 0.0002, Bootstrap), and were, everything else kept equal, more frequent after skips and less likely after correct trials. Of the four signals in the baseline, only OpticF_I_ had a positive significant association with skips (Fig. 5f; *p* = 0.0006, bootstrap), suggesting that mice are more likely to perform facial movements while they are distracted from the task. FR and Synch innovations had no explanatory power for skips (*p* = 0.19 and *p* = 0.90 for FR_I_ and Synch_I_ respectively; Bootstrap). Thus, cortical desynchronization innovations had no association with engagement for our mice (Fig. 5f).

## Discussion

Our main finding is that the effect of spontaneous cortical fluctuations on perceptual accuracy is only evident after errors, with mice making more accurate choices after errors when baseline activity was higher and more desynchronized (Fig. 3c-f). This outcome dependence could not be explained through the existence of epochs where cortical fluctuations are linked to accuracy and where errors are simultaneously more prevalent (Fig. 3h), nor through the presence of a particular baseline state favorable for accuracy found only after errors (Fig. 4). Instead, errors appear to permit baseline fluctuations to become associated with choice accuracy, consistent with a gating role. Discrimination accuracy was not associated to pupil dilation or facial movement during the baseline, but these two signals did show associations with measures of responsivity. Pupil dilation predicted the ability of the mice to withhold responding during the delay period, an ability which tended to also be associated with desynchronization (although not significantly, Fig. 5d). Facial movement clearly predicted whether the mice would disengage in a particular trial (Fig. 5f) and also, to a smaller extent, premature responding (Fig. 5d), whereas baseline neural activity did not predict engagement (Fig. 5f).

There is renewed awareness^30,31^ that observed covariations between neural activity and behavior might reflect the association of these covariates with slow unmeasured processes^28^. Indeed, the physiological and behavioral signals we analyzed all displayed slow trends of variation across the recording session (Figs. 3-5, Supplementary Fig. 2a) as well as auto- and cross-correlations spanning several trials (Supplementary Fig. 2b). In addition, pupil size is clearly affected by licking and reward in the previous trial^37^ (Fig. 4). To prevent these slow trends from contributing to our results, our approach consisted in regressing variables of interest not on the raw measured baseline signals, but on their residuals from predictive models based on past information, which we refer to as innovations (Fig. 2, Supplementary Fig. 3). The use of innovations reversed the sign of the association between PupilS and Synch, and surprisingly revealed that PupilS is exclusively associated to Synch and not to FR – the latter being only a function of movement in the baseline (Fig. 2a,c). Although baseline innovations are uncorrelated in time (Fig. 2c), the behavior of the mice still displays slow fluctuations and trends (we did not apply the same whitening procedure on behavioral measures to retain their categorical character). Several lines of evidence argue against the presence of these slow components in behavior playing a role in our findings. First, one would not expect them to lead to associations with our baseline innovations since innovations don’t have any slow components. In addition, we modeled slow behavioral trends explicitly through the use of the session trend regressor TrN (which generally has explanatory power, Figs. 3-5), in order for TrN to ‘explain away’ any correlations between behavior and marginal slow components in the innovations. For instance, TrN is effective at explaining away a correlation between accuracy (unconditional on outcome) and raw baseline FR (data not shown). Second, baseline innovations do not explain accuracy unless one conditions on outcome (Fig. 3a). And third, if the correlations between baseline neural fluctuations and accuracy after errors (Fig. 3c) was due to the existence of slow fluctuations, it would be also present when conditioning on the outcome of the next (as opposed to previous) trial, a prediction which we tested, but which was not consistent with our data (Fig. 3h). We conclude that the correlations we reveal between baseline fluctuations and behavior are indeed the reflection of true associations between these signals with a time-scale of a single trial. Early studies emphasized changes in brain state associated to distinct behavioral modes, such as exploration or quiescence^3,6,7^. More recent work has highlighted faster modulations of brain state within a single behavioral context^8,9,42^. Our results add another layer to this characterization, by showing that the *functional role* of cortical state is under the control of performance monitoring brain systems. The fact that these results were obtained using innovations demonstrates that the dynamics of this form of control are rapid, and can pop in and out within a single trial.

Our results suggest performance monitoring systems shape the effect of brain state in sensory cortex. A recent study characterized the role of a projection from the anterior cingulate cortex (ACC) to the visual cortex (VC) on performance monitoring in the mouse, showing that post-error increases in performance in a visual attention task can be mediated by this projection^43^. Although the authors did not interpret their findings in the context of modulations in cortical state, there are interesting parallels between their results and our findings. Optical pulsatile activation (30 Hz) of the ACC to VC projection resulted in decreases in low-frequency LFP power (consistent with a decrease in our Synch measure, Fig. 1h,i) and increases in high-frequency power (consistent with an increase in FR^44,45^) in the visual cortex – akin to our favorable state for accuracy after errors. Interestingly, a behavioral effect of either excitation or suppression of this projection was only observed when the manipulation was performed in the baseline period after errors. These findings suggest that the favorable state for accuracy after errors we identified might signal the successful recruitment of performance monitoring frontal networks. More generally, both studies suggest that neither particular baseline cortical states nor the activation of top-down projections from performance monitoring structures onto sensory areas are in themselves sufficient for behavioral performance to be enhanced. Rather, they seem to be a part of a more global set of brain processes (whose full extent remains to be elucidated) triggered by errors.

Another recent study by Jacobs et al.^21^ examined similar questions as our study in a visual discrimination task, assessing cortical fluctuations using widefield imaging. Our results are consistent with theirs regarding the lack of effect of cortical state on accuracy when trial outcome is not considered (Fig. 3a), but the outcome-dependent relationship between cortical state fluctuations and accuracy which we revealed was not addressed in this study. Jacobs et al., however, found cortical desynchronization to to be associated with engagement, whereas we did not (Fig. 5f). This discrepancy could be related to overall differences in skip probability across the two studies or to task differences such as the existence of a delay period in our study, or the absence of self-initiation in our task versus self-initiation by lack of movement in Jacobs et al. We did, however, find associations between baseline innovations and responsivity when examining premature responses, which, ceteris paribus, tended to be preceded by pupil contractions and synchronized baseline activity (Fig. 5d). This is interesting given that, in tasks without a delay period, it is pupil dilation^10^ and desynchronization^21^ that are associated with faster RTs and ‘false alarms’. On the other hand, the result is expected given the well known association between pupil dilation and inhibitory control^40^. In an anti-saccade task, for instance, it was found that pupil size was bigger before correct anti-saccades than before incorrect pro-saccades in anti-saccade trials^39^. That a diversity of cognitive processes converge on pupil dilation is consistent with its dependence on different neuromodulatory systems^37,46,47^. In tasks with a delay period, explanatory accounts of pupil dilation based on distractability or exploration^48,49^ and cognitive control^4,40^ appear to make opposite predictions regarding responsivity. In our task, processes associated with control seem to have a stronger hold on the pupil signal.

Our results, together with those from previous studies^21^, demonstrate that high-level discrimination performance can be sustained relatively independently of cortical synchronization (in our case after correct trials). What general conclusions can be derived from these findings regarding the relationship between cortical state and sensory discrimination accuracy? In addressing this question, we first note that, in humans, good levels of performance can be obtained in well rehearsed tasks, with high degrees of automaticity and in the presence of frequent feedback – exactly the conditions present in psychophysical tasks like ours – in the absence of the kind of mental effort associated with focused attention^50,51^. These ‘flow’ states, in which subjects experience dissociation and lack of self-consciousness, are thought to arise when skills and demand are matched^52^. Interestingly, brain structures implicated in performance monitoring and engaged by task errors, such as the ACC and medial prefrontal cortex^43,53,54^, are downregulated during flow^55,56^. We hypothesize that, when discriminating simple sensory stimuli, rodents (especially if not forced to perform at psychophysical threshold) might operate in a state equivalent to ‘flow’ during streaks of correct trials, and that during these states the involvement of sensory cortex in discriminative choice might be low. This outcome-dependent reconfiguration of brain pathways supporting choice would explain why, after correct trials, accuracy is independent of cortical state fluctuations in our task and of the activation of top-down inputs to the visual cortex in the study of Norman et al.^43^Errors might promote cortical involvement, which would render cortical spontaneous fluctuations relevant for choice, both directly through their potential effect on sensory representations^14–18^, and also indirectly as markers of focused attention^43,57,58^.

Overall, our findings suggest that the relationship between spontaneous activity fluctuations in sensory cortex and choice performance is labile and context-dependent, and raise the possibility that the brain systems supporting such performance might display equivalent flexibility.

## STAR Methods

### Resource availability

#### Lead Contact

Further information and requests for resources and reagents may be directed to, and will be fulfilled by the lead contacts, Davide Reato and Alfonso Renart.

#### Materials Availability

This study did not generate new unique reagents. All reagents are commercially available.

#### Data and Code Availability

All data and custom MATLAB scripts used to analyze the data are available upon reasonable request.

### Experimental model and subject details

All procedures were reviewed and approved by the Champalimaud Centre for the Unknown animal welfare committee and approved by the Portuguese Direção Geral de Veterinária (Ref. No. 6090421/000/000/2019). All experiments were performed using male C57BL/6J mice that were housed on a 12h inverted light/dark cycle.

## Methods details

### Head bar surgery

During induction of anaesthesia, animals (6-8 weeks of age, 20-22 g body weight) were anesthetized with 2-3% (volume in O_2_) isoflurane (Vetflurane, Virbac) and subsequently mounted in a stereotactic apparatus (RWD Life Science) on a heating pad (Beurer). Once animals were stably mounted, isoflurane levels were lowered to 1-1.5% and the eyes were covered with ointment (Bepanthen, Bayer Vital). The head was shaved and the scalp cleaned with betadine. A midline incision was performed to expose lambda and bregma, which were subsequently used to align the skull with the horizontal plane of the stereotactic frame by measuring their position with a thin glass capillary (Drummond Scientific). The skull anterior of bregma was exposed by cutting a small area of skin. The exposed area was cleaned with betadine and slightly roughened by scraping it with a surgical blade (Swann-Morton). Subsequently, the skull was dried with sterile cotton swabs and covered with a thin layer of super glue (UHU). To further increase long-term stability, four 0.9 mm stainless steel base screws (Antrin Miniature Specialties) were placed in the skull. The exposed skull and base screws were then covered with dental cement (Tap2000, Kerr). A custom designed head bar (22×4×1 mm, aluminium, GravoPlot) was lowered into the dental cement while still viscous until the head bar was in contact with the base screws. Subsequently, an extra drop of dental cement was applied to the centre of the head bar in order to fully engulf its medial part. The remaining skin incision along the midline was then sutured. The animals were injected with buprenorphine (opioid analgesic, 0.05 mg/kg) into the intraperitoneal cavity and allowed to recover for 3-5 days.

### Training

We adapted previously described procedures for training head-fixed mice in psychophysical tasks^59^. After recovery from head bar implantation the animals were water deprived for 12 h prior to the first handling session. In handling sessions mice were accustomed to the experimenter and being placed in an aluminium tube to restrain their movement. In the first days of handling, the tube was placed in the animal’s home cage. Once the mouse voluntarily entered the tube, it was presented with water delivered manually from a syringe at the end of the tube. This procedure therefore roughly mimicked the water delivery system in the training apparatus. Mice were allowed to drink a max of 1.5 ml of water during each handling session (30 min). Once mice were accustomed to receiving water in the aluminium tube and being handled by the experimenter, they were placed in the behavioral setup and head fixed with the two water delivery spouts approximately 1 cm in front of their mouth. To adapt them to head fixation, free water was delivered upon licking at either of the water delivery spouts. Lick detection was based on junction potential measures between the aluminium restraining tube and the stainless steel lick spout^60^. After triggering and consuming 15 rewards (single reward size: 3 *µ*l), training proceeded as follows.

In the first stage of training, every 3.1 s a random high (distribution: 22-40 kHz, presented sound randomly selected each trial, category threshold at 14 kHz) or low (distribution: 5-8.5 kHz) frequency sound was presented to both ears at 60 dB SPL for 600 ms, indicating at which of the spouts water was available (mapping is counterbalanced across animals). 150 ms after sound onset, a green LED flash of 50 ms indicated the onset of the 1.5 s response period. If the first lick in the response period occurred at the correct water spout, a 3 *µ*l water reward was delivered. In order to facilitate the animals’ engagement, a free water drop was delivered 150 ms after sound onset in a random 10% subset of trials. Once mice were readily trying to trigger water rewards by licking at either lick spout after sound presentation (minimum of 18 out of the last 20 trials without free water), the sound duration was reduced to 150 ms, followed by a 1s response period. The inter-trial interval (ITI) was drawn randomly from a set of four possible values: 3, 4, 5 and 6 seconds. After mice were engaged in the new timing of the task, free water delivery ceased and incorrect responses were punished by an additional 6 s time delay in between sound presentations. As soon as the animals had learned to correctly respond by licking at the appropriate water delivery spout in at least 34 of 40 consecutive trials, the response delay was introduced, by gradually delaying the appearance of the go signal. Impatient licks triggered the abort of the trial and were signalled with white light flashes. The delay period was increased in 10 ms increments as long as the animal performed at an accuracy of at least 80% for maximal five increments per session.

In the following weeks of training the difficulty of the presented frequencies was gradually increased by approximating the range of possible low and high frequencies. These increases were performed in 19 increments, depending on a low bias and and high performance (bias ≤ 20%; performance ≥ 80%, only one change per training session), until a final frequency distribution of low (9.9-13 kHz) and high (15-20 kHz) frequencies was reached. After reaching the final frequency distributions, mice were presented with three fixed frequencies per condition (Low: 9.9, 12 and 13 kHz; High: 15, 16.3 and 20 kHz), with the easy conditions presented only 15% of the times to obtain more error trials and hard trials presented in 8% of the trials. Due to the resulting low number of hard trials per behavioral session, their presentation was omitted during most acute recording sessions (24 of 36). Excessive bias or disengagement at any time during the training were corrected by delivering free water at the unpreferred spout right after stimulus presentation until the animal readily responded again. All such intervention trials and the trials subsequent to each of them were excluded from analysis.

### Electropysysiological recordings

Six to 12 h prior to the first probe insertion in each hemisphere, mice were deeply anesthetized with 2-3% (volume in O_2_) isoflurane, mounted in a stereotactic apparatus and kept on a thermal blanket. The eyes were covered with ointment. Isoflurane levels were subsequently lowered to 1-1.5%. The animals head was placed in a stereotactic frame using the head bar. The skin covering the areas above the recording sites and the midline was removed and the exposed skull was cleaned from periostium with a surgical scalpel blade and cleaned with betadine and dried with sterile cotton swabs. Subsequently a small craniotomy was performed above the desired recording site (2.8 mm posterior, 2.2 mm medio-lateral to bregma under a 35° medio-lateral angle). The exposed dura mater was opened using a small needle (BD Microlance^*TM*^ 0.3 × 13 mm) and subsequently the recording silicone probe (BuzA64sp, Neuronexus) was slowly lowered to the desired depth (2.6 mm from brain surface). Probes were inserted with the shanks in a medio-lateral orientation, so that the six shanks in the final position approximately span the cortical layers (Supplementary Fig. 1a). Neural activity was digitised with an 64 channel headstage (Intan) at 16 bit and stored for offline processing using an Open Ephys acquisition board (Open Ephys) at a 30 kHz sampling rate. Behavioral sessions and storage of recording neural signals started only 10-20 min after probe insertion to allow for tissue relaxation and stabilization of the recording. Recording sessions were limited to three recordings per hemisphere in each animal due to the tissue damage caused by probe insertion. In the final recording session in each hemisphere the probe was coated with DiI (Vybrant*™* DiI, Invitrogen) to confirm correct placement of the recording probe histologically.

### Data Set

We recorded neural activity in 36 behavioral sessions from 6 mice (3 recordings per hemisphere). Out of these, 23 sessions had at least 100 trials and a behavioral sensitivity (*d*′ from signal detection theory) of at least 1, and were considered for further analysis. We confirmed histologically that recordings were made in both primary as well as ventral and dorsal auditory cortex (see Supplementary Fig. 1 for an example recording). Behavioral sessions from one mouse (3 sessions) were discarded from the dataset because it was not possible to properly estimate the size of its pupil due to eyelid inflammation. Inclusion of this mouse does not affect aggregate results regarding the effect of cortical desynchronization. Unless otherwise specified in the text, we didn’t consider for analysis the first 10 trials in each behavioral session during which the mice are adjusting to the setup and the position of the licking ports is being fine tuned. We also did not consider trials where the current or the previous trial were free rewards (trials in which the experimenter delivered a free reward to re-engage the animal). For the analysis on accuracy we considered the first lick within the response window (0.7 to 1.7 s after sound onset) which was also used to determine if animals would be rewarded. For the analysis of engagement, ‘skips’ were defined as trials in which no licks were detected in the first 2 s since stimulus onset. Premature responses were defined as trials in which the first lick occurred before the go-signal, at 0.65 sec.

## Quantification and statistical analysis

### Videos recording and analysis

We collected videos of mice performing the task at 60 fps using regular USB cameras without an infrared (IR) filter and applying direct IR illumination to increase pupil contrast (Supplementary Fig. 1b). From the videos, we extracted a proxy for face movement and one for arousal. For face movement, for each recording session, we selected a ROI around the face of the animal and computed the average magnitude of the optic flow in that ROI (using Lucas-Kanade method^61^). To compare across sessions, we z-scored the optic flow session by session. What is referred to as OpticF in the text corresponds to the median OF in the baseline period (2s before stimulus presentation). As a proxy for arousal, we estimated pupil size. We used DeepLabCut^62^(DLC) to detect points around the pupil frame by frame and then estimated the pupil size as the major axis of an ellipse fitted using those points (for robustness of the pupil estimates, we further smoothed the data by applying a robust local regression using weighted linear least squares and a 1^*st*^ degree polynomial model with a 250 ms window – `rlowess` in MATLAB). For training the model using DeepLabCut, we labelled 8 points in 20 frames for each of the 20 behavioral sessions. To remove frames where the detection was poor, we only considered those where the average likelihood of the DLC detection was higher than a threshold (0.8). Finally, for each session, we normalized the pupil by the 2% lowest values in the session (so, for example, 100% means a 100% increase in pupil size relative to its smallest values). What we referred in the main text as PupilS represents the median values of the pupil in the baseline period (2 s before stimulus presentation).

### Spike Sorting

Spike events were detected using Kilosort2^63^ (github.com/MouseLand/Kilosort2) and subsequently manual clustering was performed using phy2 (github.com/cortex-lab/phy) to remove artifact clusters. We didn’t use unit identity in any of our analyses, which pertained only to the structure of the population (‘multiunit’ MUA) activity.

### Estimation of baseline firing rate and synchrony

We described baseline neural activity in each trial using two variables, the population firing rate (FR) and synchrony (Synch; Fig. 1f). We estimated FR as the average number of spikes of the multiunit activity (MUA) in the baseline period (average in time and across the number of units). To estimate synchrony, we first pooled all spikes from the units in the baseline period. Then computed the magnitude of the standard deviation across time of the instantaneous firing rate (in bins of 20 ms) of the population (which is a measure of the population averaged covariance between all pairs^26^) and divided it by the average of the same quantity calculated for 100 surrogates where the spike times of the MUA in that particular baseline are randomly shuffled (Fig. 1e). We used this measure because we observed that it is less dependent on overall number of spikes in the baseline period than related measures such as the coefficient of variation of the MUA across time^18,26^. This measure of synchrony is ‘normalized’, with a reference value of 1 expected if the neural population is asynchronous and neurons fire with Poisson-like statistics. In Fig. 1g, we assessed synchrony using the coefficient of variation (CV) of the MUA, defined as the ratio between the standard deviation and the mean the spike count of the MUA across each 20 ms period in the baseline period.

### Spectral analysis

We performed spectral analysis using the Chronux Matlab Package (http://chronux.org). In particular, we used the function `mtspectrumpt.m`, which uses a multitaper approach to calculate efficiently the power spectrum of a point process. In Fig. 1h, for each of the four example baseline periods, we used a value for the time-bandwidth parameter *TW* = 10. For Fig. 1i, since additional smoothing is provided by the average across trials, we used *TW* = 5. In each case, we used the recommended 2*TW* – 1 tapers to calculate the spectrum in each baseline period. Each power spectrum was normalized by the mean power for all frequencies above a high frequency cutoff of 10 kHz (the sampling rate of the recordings was 30 kHz), which is equivalent to a normalization by the firing rate within that baseline period (since the high frequency limit of the spectrum of a point process is the firing rate).

### Innovations

We ‘cross-whitened’ the four signals under analysis (FR, Synch, OpticF and PupilS) by making linear fits of each of them separately for each session, using as regressors the outcome in the previous 10 trials (1: reward; 0: no reward), the values of four signals in the previous 10 trials and the current trial number (TrN, to account for within sessions trends). Each regression thus specified 51 parameters plus the offset. We then defined the innovations FR_I_, Synch_I_, PupilS_I_ and OpticF_I_ as the residuals of this linear fit (Supplementary Fig. 3). In Fig. 4 we address the influence of the outcome of the previous trial on the four baseline innovations. We did this by trying to explain previous outcome using a GLMM based on these signals plus the session trend. For this fit, the innovations were modified by excluding previous-trial outcome as a regressor (since their relationship to outcome is the target of the analysis).

### Generalized linear mixed models (GLMM)

To analyze the behavioral and neural data we used generalized linear mixed models^64^ (GLMM, using the function `fitglme` in MATLAB) using recording session as a random effect for both slopes and offset. When fitting continuous variables (e.g., FR_I_ in Fig. 2d) we used a linear mixed model. When fitting binary variables (such as accuracy or skips) we used a binomial distribution and a logit link function. In order to prevent global covariations between session-by-session differences in the marginal statistics of the predictors and the prediction targets to contribute to the trial-by-trial associations that we seek to reveal, we always z-scored all predictors separately *within each session*. In all fits, we also include a regressor with the number of the trial in the session (TrN) to account for session trends in the target of the fit. In Fig. 3c,e, we evaluated the *joint* effect of FR_I_ and Synch_I_ on choice (rightmost predictor). To do this, we first constructed a joint predictor by projecting each z-scored (FR_I_(z), Synch_I_(z)) pair (for each trial) onto an axis with -45 deg slope for Fig. 3c (so that the joint predictor would take large positive values when the baseline state was favorable after errors), or 45 deg slope for Fig. 3e (using the same reasoning after corrects). We then run GLMMs in which the two separate FR_I_ and Synch_I_ predictors were replaced by the single joint one. In Fig. 3c,e, we only show the value of the joint coefficient in these new GLMM fits. The values of all other predictors were not different.

### Statistics

We estimated the uncertainty of the estimates of the coefficients of our GLMM fits using bootstrap resampling^32^. We resampled with replacement ‘hierarchically’, so that the number of trials from each recording was preserved in each global surrogate. Distributions of the magnitude of each coefficient and associated 95% confidence intervals (CI) came from 5000 resamples. In figures, we always display median, interquartile range and 95% CI for each coefficient. *p* values for the null hypothesis of a coefficient being equal to zero were computed using the quantile method^32^, i.e., twice the value of the fraction of resamples with opposite sign as the estimate of the coefficient from the data. For consistency, we verified that estimates of significance obtained using bootstrap confidence intervals for parameters agreed with parametric estimates from `fitglme` (Supplementary Fig. 4) which uses an approximation to the conditional mean squared error of prediction (CMSEP) method^65^. To test for differences in accuracy after a correct versus an error trial (Fig. 4b), we computed, for each recording, the difference between the median accuracy of trials where the previous trial was correct and the median accuracy of trials where the previous trial was an error. We assessed the significance of this difference using a Wilcoxon signed rank test. To fit psychometrics curves, we used the psignifit MATLAB toolbox^66^(github.com/wichmann-lab/psignifit/). When fitting an aggregate psychometric across sessions, we weighted each trial by the proportion of trials its corresponding session contributes to the whole dataset. To test for differences in the slope of the psychometric functions in Fig. 4d,f conditional on whether the baseline state was favorable or unfavorable, we used the difference in slope between fits of the aggregate data conditional on the state of the baseline as a test statistic. To assess the significance of this difference, we first computed the distribution of the test statistic under a null hypothesis of no difference implemented by randomly shuffling, within each session separately, the label that signals whether the baseline for a trial is favorable or unfavorable, and we then computed the fraction of the surrogates from this distribution for which the value of the test statistic was equal or larger than the in the actual observed data. Unless otherwise stated, data across recordings is reported as median ± MAD (median absolute deviation).

## Acknowledgements

We thank Julien Fiorilli for help developing the lick detection hardware, the Vivarium and Hardware scientific platforms at Champalimaud Research for support, and Leopoldo Petreanu, Michael Orger, Jaime de la Rocha and Tiffany Oña for comments on the manuscript. D.R was supported by a Fundação para a Ciência e Tecnologia postdoctoral fellowship (SFRH/BPD/119737/2016) and a Marie Skłodowska-Curie postdoctoral fellowship (H2020-MSCA-IF-2016 753819), R.S was supported by a doctoral fellowships from the Fundação para a Ciência e a Tecnologia. AR was supported by the Champalimaud Foundation, a Marie Curie Career Integration Grant PCIG11-GA-2012-322339, the HFSP Young Investigator Award RGY0089 and the EU FP7 grant ICT-2011-9-600925 (NeuroSeeker).

## Author contributions

D.R., R.S. and A.R. conceived the project. R.S conducted the experiments with support from A.T.M. D.R. and A.R. designed the analyses. D.R. conducted the analyses and curated the data. A.R. wrote the manuscript with feedback from all authors.

## Supplemental information

**Supplementary Fig 1.**
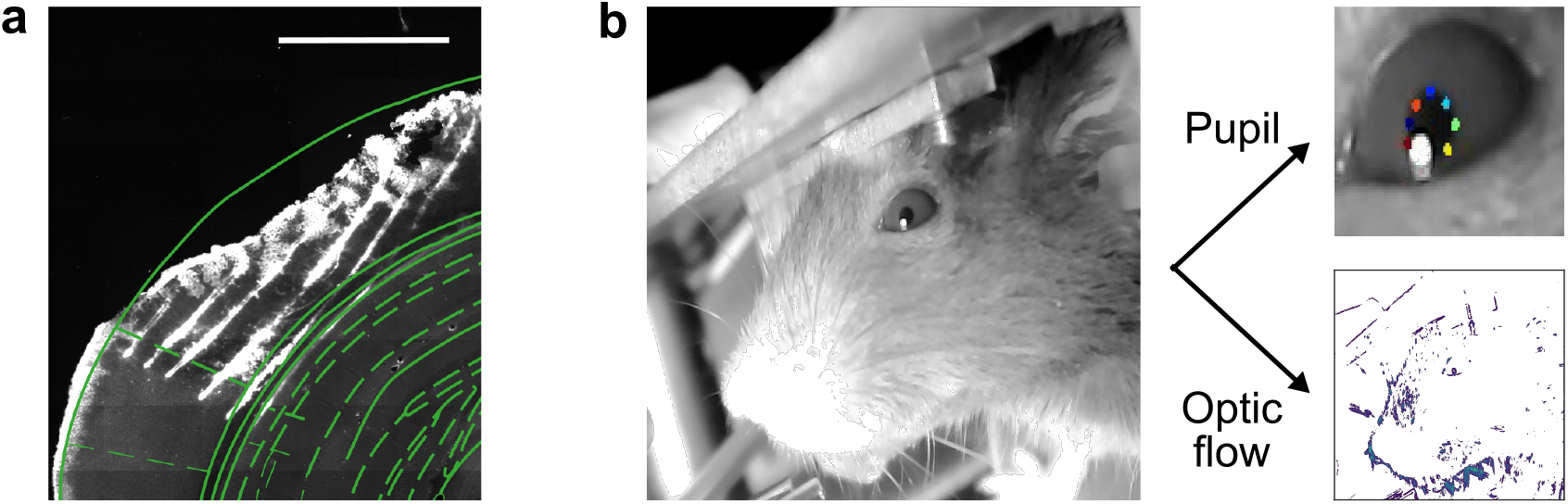
Histology and video analysis, related to Fig 1. **(a)**. Reconstruction of a brain slice with the shanks of the probe marked with DiI. Shank tips in this recording were in the primary auditory cortex (areas adapted from the Paxinos brain atlas). **(b)** Left. Image of the face of the mouse in our set up. Right. From these videos we extract pupil size using DeepLabCut^62^, marking 8 points to characterize the ellipse for each pupil (top) and optic flow – to quantify facial movement – by counting the number of pixels (in color in the figure) which change value across adjacent frames.

**Supplementary Fig. 2.**
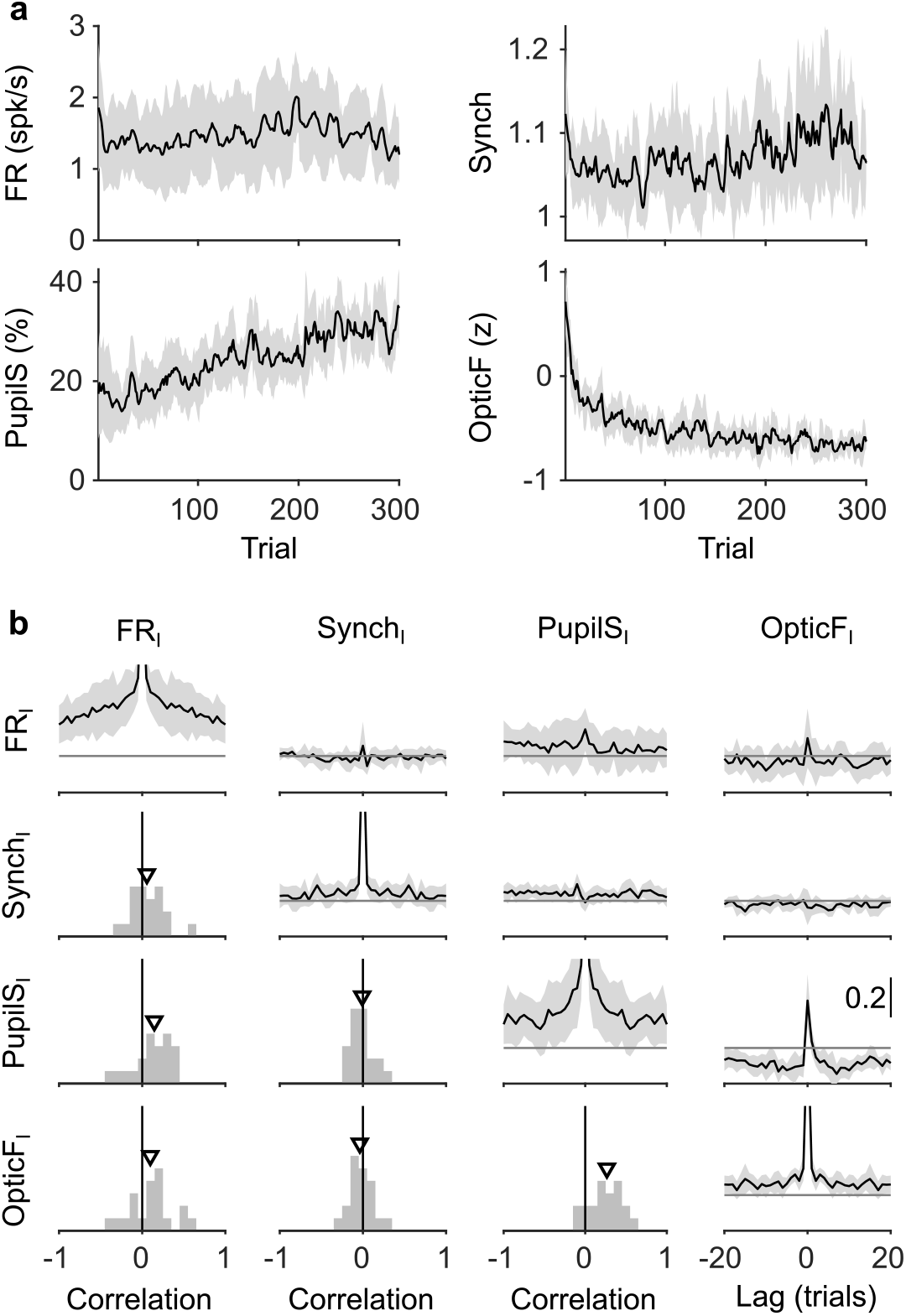
Slow trends of baseline signals during the session, related to Fig 2. **(a)** Median (black) and MAD (gray shading) across sessions of each of the four baseline signals. All the signals display slow trends and in some cases monotonic increases or decreases through the recording session. **(b)** Diagonal and above shows the auto- and cross-correlations of each of the four baseline signals (median ± MAD). Below diagonal shows the histogram of the instantaneous correlation between each pair of signals across sessions. Triangle is the median across sessions (same format as Fig. 2c). Non-zero values of auto- and cross-correlations far from zero lag reflect existence of slow time-scales, which are eliminated by our cross-whitening procedure (Methods, Supplementary Fig. 3) and are thus absent from the equivalent analysis performed on innovations (Fig. 2 in the main text.)

**Supplementary Fig. 3.**
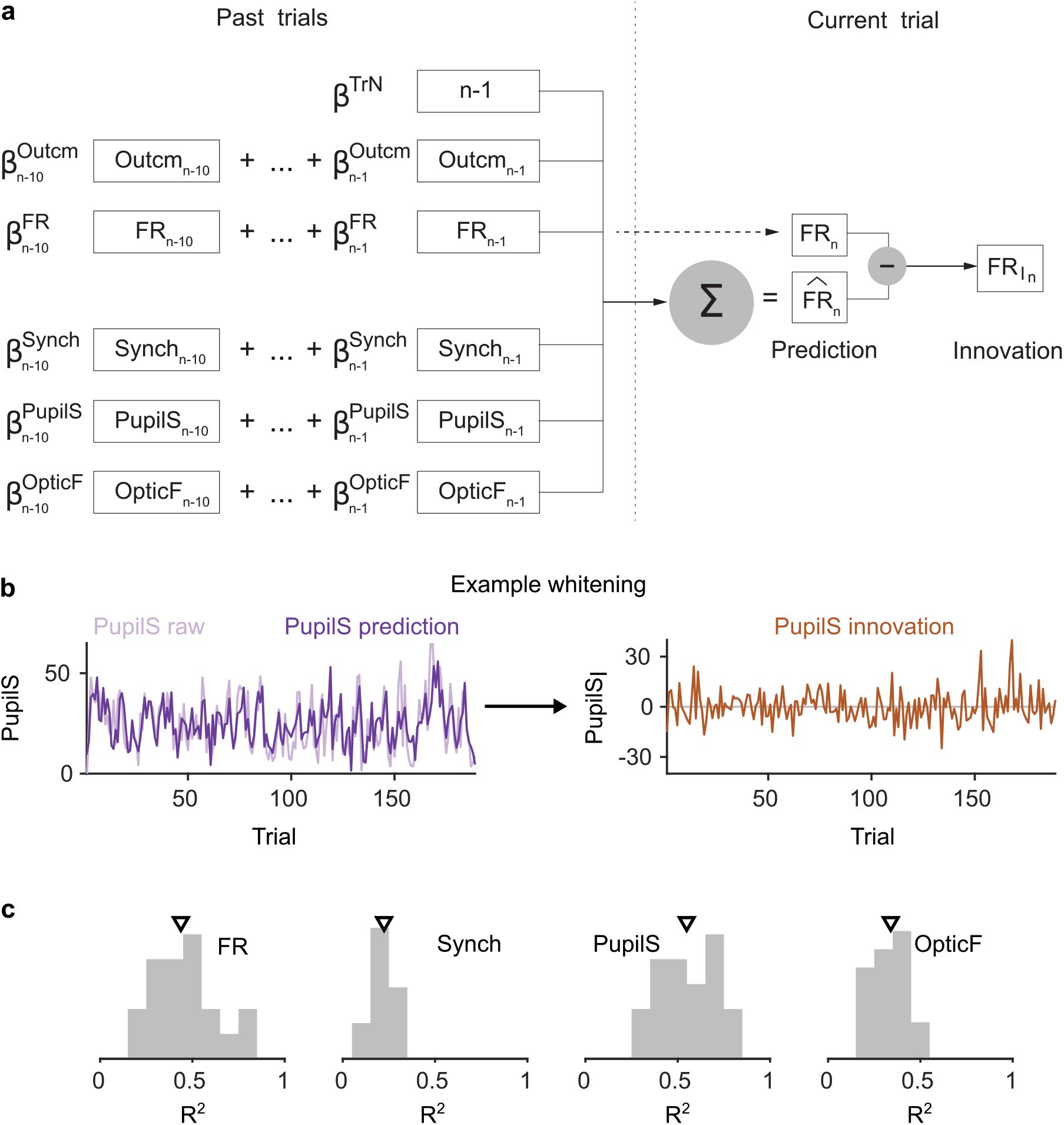
Constructing innovations by cross-whitening, related to Fig 2. **(a)** Schematic description of the linear fit and associated residuals used to generate the FR innovations. The same procedure was used for Synch, PupilS and OpticF. **(b)** Example traces for obtaining the PupilS innovations in one recording. Left, raw and linear fit of the PupilS. Right, residuals. **(c)** Histogram across sessions of the fraction of variance (*R*^2^) explained by the linear fits of each of the four signals. Triangle is the median of each histogram.

**Supplementary Fig. 4.**
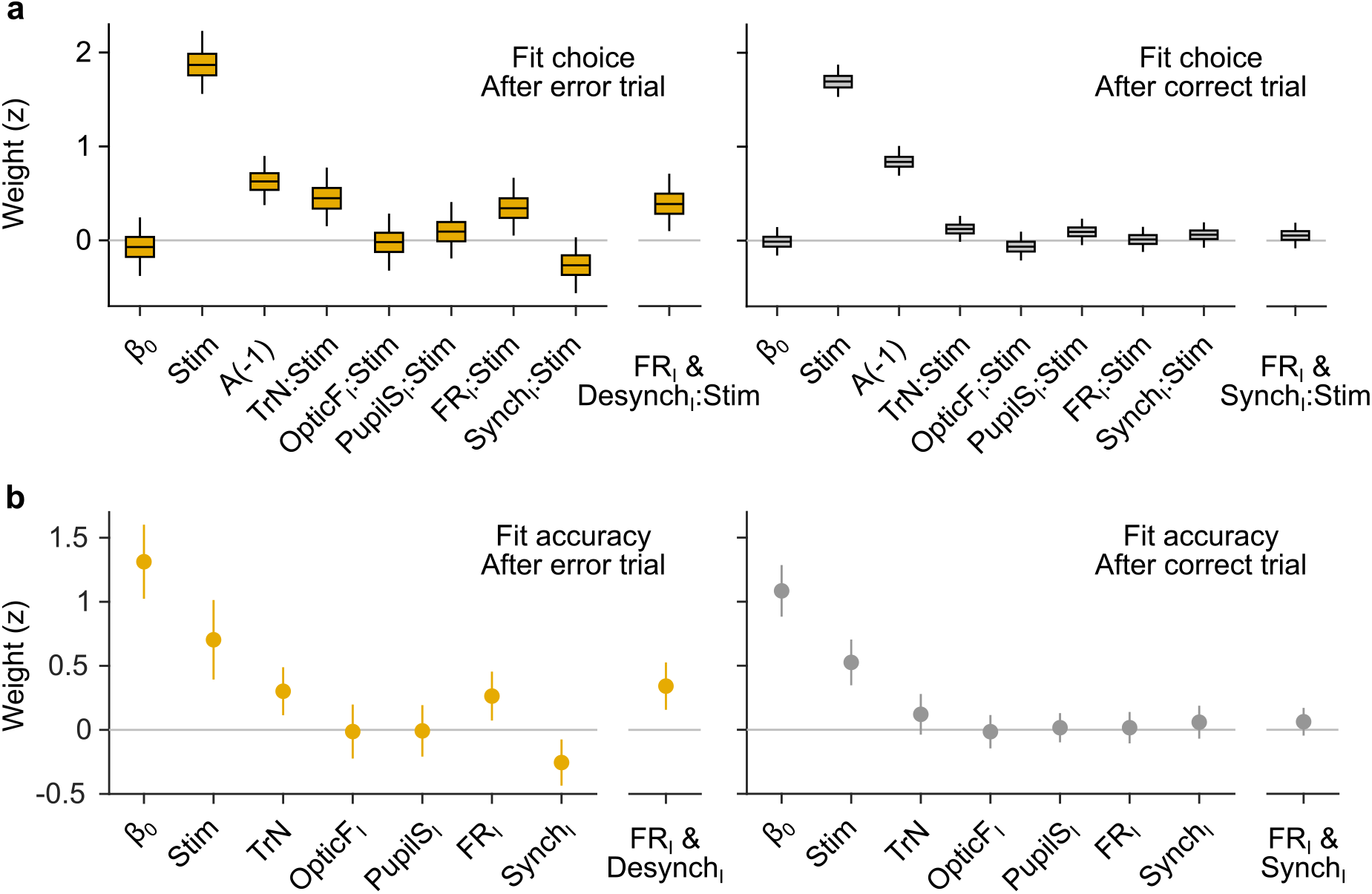
Robustness of the association between brain state and accuracy, related to Fig 3. **(a)** Coefficients of a GLMM fit for choice after error (Left) and correct (Right) trials (as opposed to accuracy in Fig. 3c,e). We use the same predictors, except that now the history predictor reflects previous choice (as opposed to previous outcome), and that the remaining predictors are included as interaction terms with the stimulus predictor. The coefficient for previous choice is positive (*p* < 0.0002) signaling that the mice tend to repeat their previous choice (to a slightly smaller extent after an error trial). Despite reflecting interaction terms and not main effects, coefficients for innovations are qualitatively similar to their counterpart in Fig. 4, although the association of choice accuracy with baseline fluctuations is slightly weaker (95% CI for the synchrony innovation now touches zero; *p* = 0.08, bootstrap). The favorable state predictor (rightmost box) is still significant (*p* = 0.008, bootstrap). **(b)** Parametric estimation of confidence intervals for accuracy fits. Equivalent to Fig. 3c,e, but circle and bars show the mean and 95%CI for each coefficient reported by `fitglme` (Methods) using an approximation to the conditional mean squared error of prediction (CMSEP) method^65^.

**Supplementary Fig. 5.**
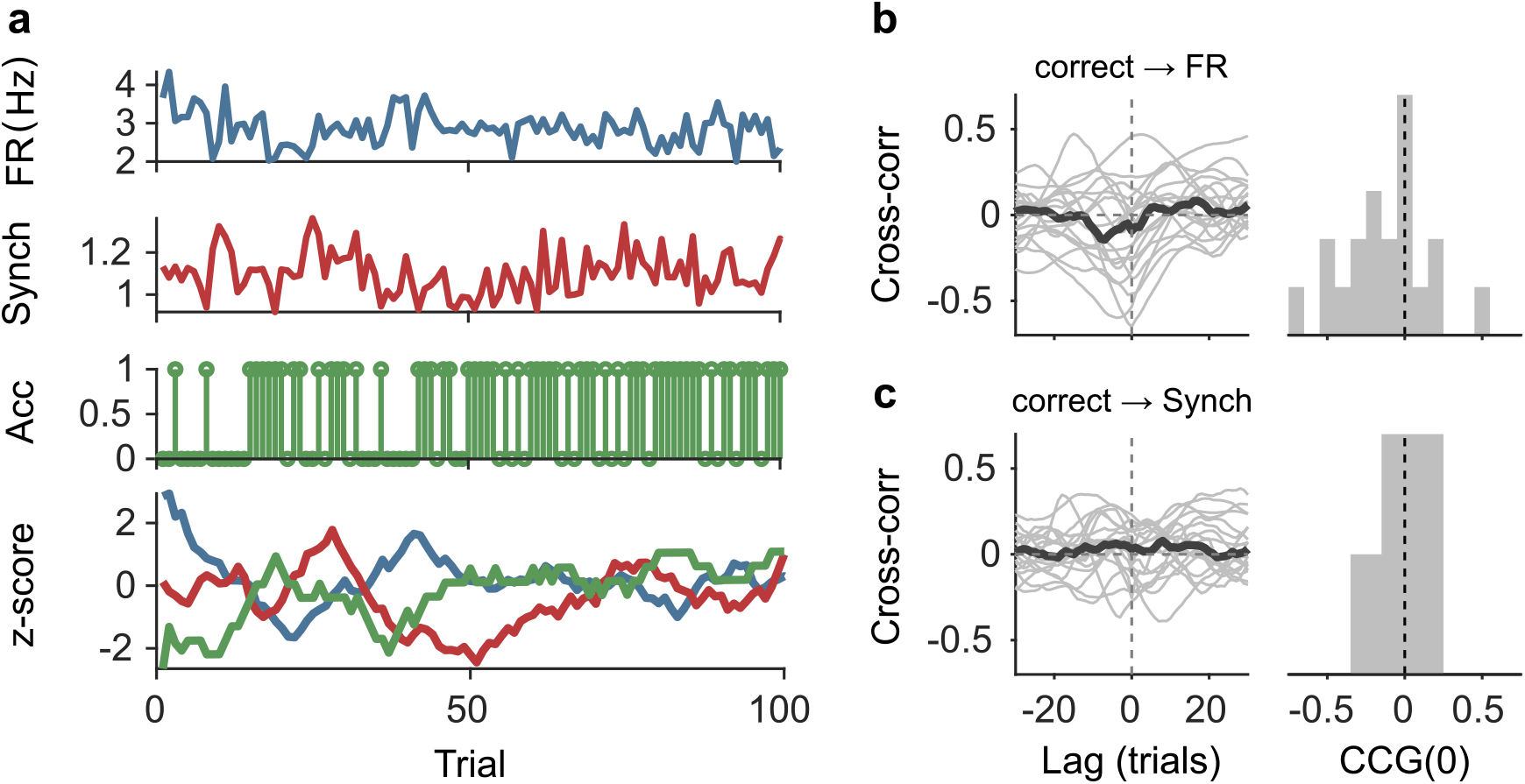
Lack of association between slow cortical state fluctuations and accuracy, related to Fig 3. **(a)** For an example session, we show the raw FR (top), Synch (middle top) during the baseline and accuracy (middle bottom) in that trial. Bottom. We smoothed each of these signals with a running window of 10 trials, removed the session wide linear trend, and z-scored. **(b)** Left. Cross-correlation function between the smoothed accuracy and FR time series. Each gray line is a recording and the black line is the mean. Right. Histogram across recordings of the cross-correlation function at zero lag. **(c)** Same as (b) but for the cross-correlation between the smoothed accuracy and Synch time series.

**Supplementary Fig. 6.**
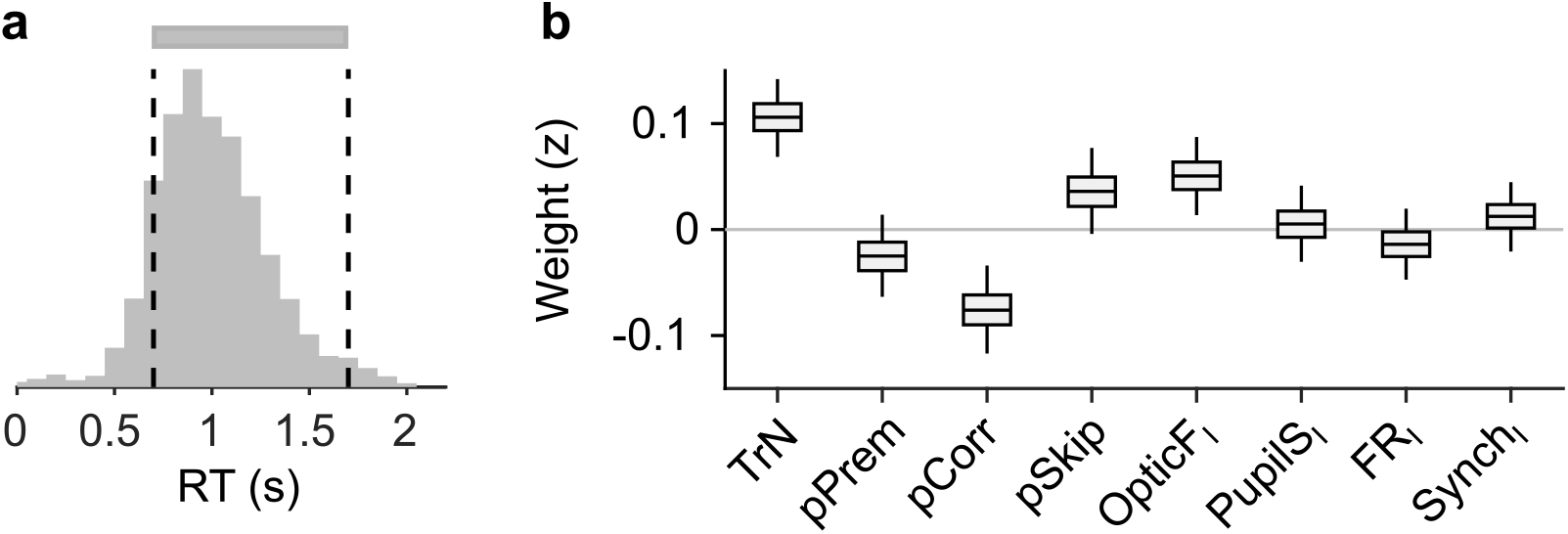
Explaining reaction time in valid trials, related to Fig 5. **(a)** Histogram of RTs (equivalent to Fig. 5a). For this figure, we attempt to explain RTs within the two dashed lines (gray bar), i.e., during valid trials. **(b)** Coefficients of a LMM explaining RT using the same predictors as in our other analyses on responsivity in Fig. 5d,f. Coefficients for session trend and previous outcome are positive and negative respectively (*p* < 0.0002 and *p* = 0.0006, bootstrap), showing that mice tend to slow down through the session – consistent with them progressively losing motivation – and also after an error – revealing that post-error slowing down is evident despite the delay period in the task. Although the pSkip coefficient is not significant (*p* = 0.075, Bootstrap), mice tend to be slower in responding after a disengaged trial, suggesting a continuity between long RTs and lack of response. This is consistent with the positive association between facial movement in the baseline (OpticF_I_) and RT (*p* = 0.01, bootstrap), which is also present in the prediction of skips (Fig. 5f). Neither pupil size, firing rate or synchrony innovations explain RT (*p* = 0.78, *p* = 0.42 and *p* = 0.45 for PupilS_I_, FR_I_ and Synch_I_ respectively, bootstrap).

**Supplementary Table 1.**
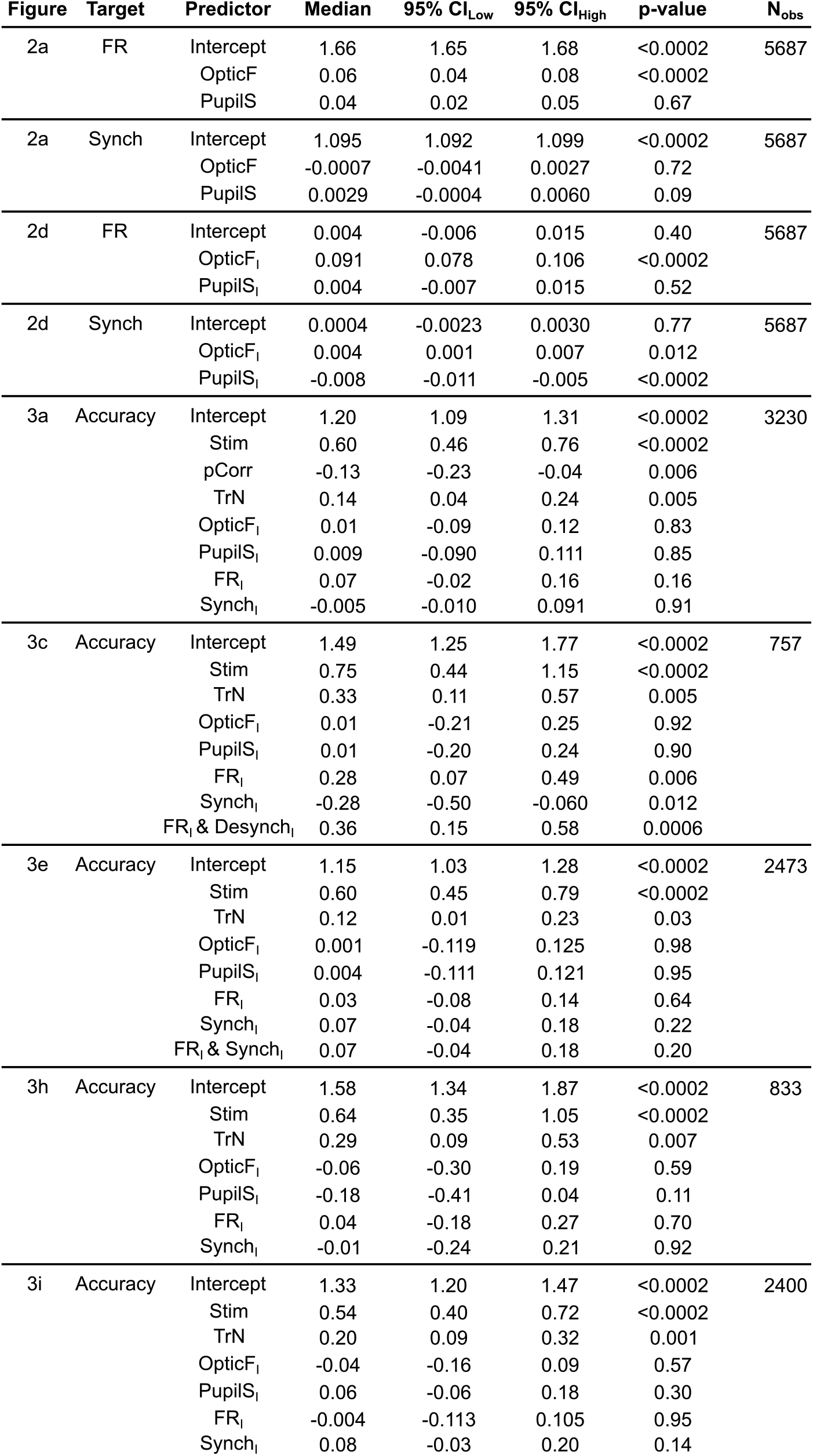
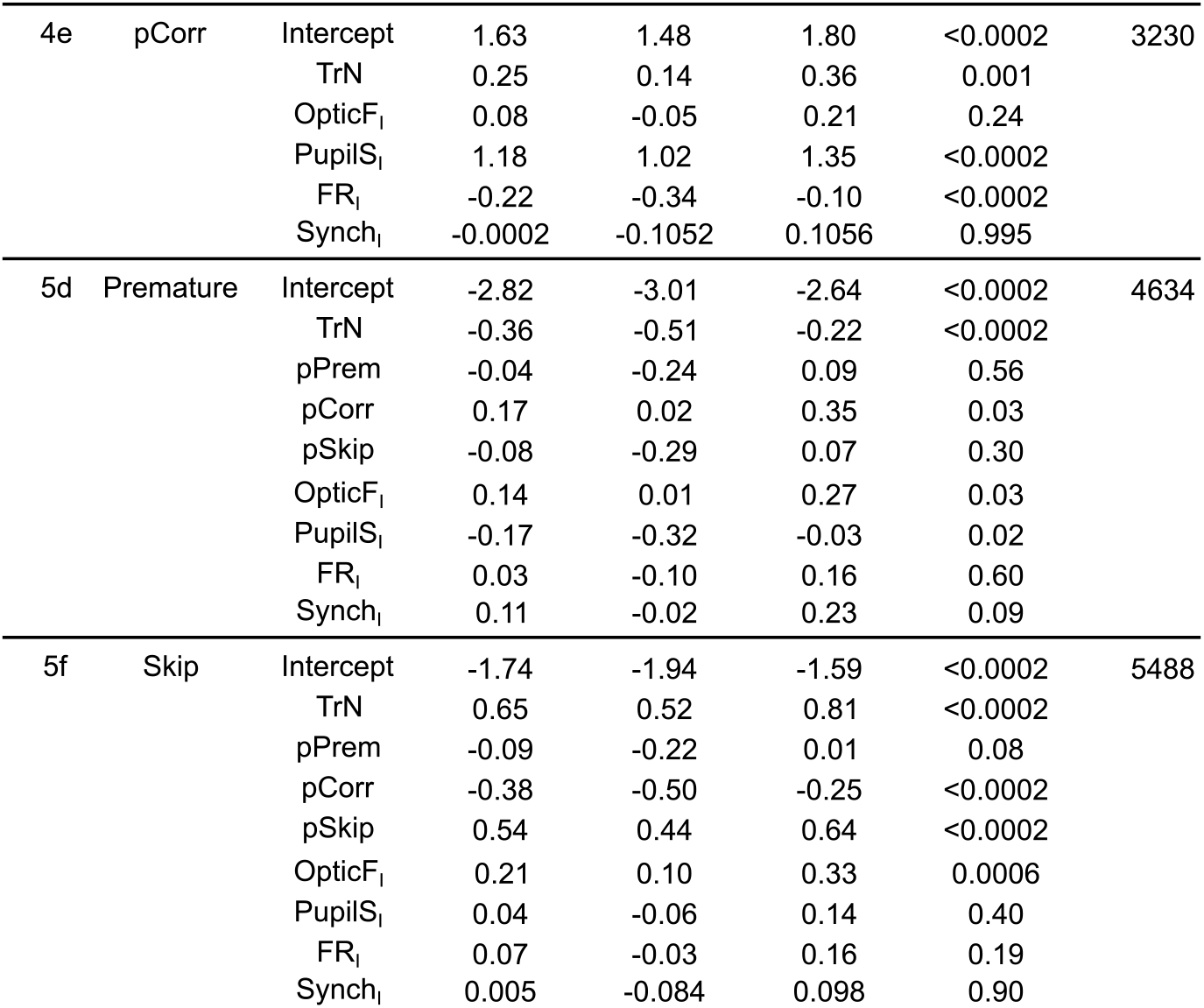
We report the statistics associated to the fixed coefficients in each of the generalized linear mixed models described in the main text. Starting from the left, each column represents: the figure in the text where the results are displayed, the prediction target, the predictors (one row per predictor), the median and lower and upper limits of the 95% confidence interval (Methods), the associated bootstrap p-Value (Methods), and the total number of observations (number of rows in the predictor matrix) in the model.

